# Verification of nanopore sequencing technology for clinical carbapenem-resistant *Enterobacterales* surveillance

**DOI:** 10.64898/2026.08.04.742736

**Authors:** Ela Sauerborn, Ebenezer Foster-Nyarko, Kathrin Schröder, Annika Sobkowiak, Samir Vargas da Fonseca Atum, Friedemann Gebhardt, Nina Wantia, Lara Urban

**Affiliations:** Helmholtz AI Institute, Helmholtz Centre Munich, Germany; Institute of Medical Microbiology, Immunology and Hygiene, Department of Preclinical Medicine, Technical University of Munich, Germany; Department of Infection Biology, London School of Hygiene & Tropical Medicine, London, United Kingdom; Institute of Hygiene, University Hospital Münster, Münster, Germany; Department of Biochemistry, Institute of Chemistry, University of São Paulo, São Paulo, Brazil; School of Medicine and Health, Technical University of Munich University Hospital, Munich, Germany; Institute for Food Safety and Hygiene, Vetsuisse Faculty, University of Zurich, Switzerland

## Abstract

Carbapenem-resistant *Enterobacterales* (CRE) pose a critical threat to global public health and often contribute to the rapid plasmid-mediated dissemination of carbapenemase genes. While established routine diagnostics can confirm the presence of the most common carbapenemases, these approaches do not resolve the genomic context of resistance and thus cannot confirm transmission events, cross-species dissemination, or atypical resistance mechanisms. Nanopore sequencing-based whole-genome sequencing (WGS) can capture this genomic context through complete *de novo* genome and plasmid assemblies. However, for routine clinical use of nanopore WGS for CRE surveillance, direct comparisons with established diagnostics and clear guidelines on required sequencing depths are needed.

We used 100 carbapenemase-producing CRE isolates from routine diagnostics at a tertiary-care hospital to compare results from WGS against established diagnostics, and determined the sequencing depth required for species identification, strain typing, carbapenemase detection, and plasmid-level epidemiology. We additionally examined 10 carbapenem-non-susceptible CRE isolates, for which routine diagnostics identified no carbapenemase gene despite phenotypic carbapenem non-susceptibility. Across all isolates, nanopore WGS reproduced routine carbapenemase family and pathogen detections, and additionally resolved the carbapenemase subtypes and their genomic context, the bacterial species and strain, and resistance mechanisms that established diagnostics had missed. Such strain typing and plasmid-level resolution are essential for infection control responses to differentiate between clonal spread of CRE, dissemination of shared plasmid, or unrelated infection events.

Our study thus strongly supports the integration of cost-efficient nanopore WGS into CRE diagnostics, surveillance, and outbreak investigation. The required sequencing depth depends on the clinical objective, with species identification being reliable at a depth of 10x, strain typing and carbapenemase detection at a depth of at least 20×, and robust plasmid-level characterisation at a depth of at least 40×. Across our CRE collection, the detected carbapenemases were mostly plasmid-borne, and predominantly encoded by relatively conserved IncN and more heterogenous IncL/M plasmids.

**Importance:** Carbapenem-resistant bacteria are among the most serious threats in modern medicine, leaving clinicians with few treatment options. Nanopore sequencing can be a powerful tool to rapidly and precisely track resistance and guide infection control, but limited comparisons with established diagnostics and uncertainty about how much sequencing data is needed currently limit routine clinical use. We show that nanopore sequencing detects all relevant carbapenemase genes identified by routine diagnostics, resolves carbapenem resistance mechanisms that standard tests miss, and increases the resolution of pathogen and gene characterizations for transmission and outbreak tracing. We provide guidance on the sequencing depth required for diagnostic tasks, from identifying species to tracking plasmid-borne resistance genes across time and bacterial species. By benchmarking nanopore sequencing against established diagnostics and matching sequencing effort to the clinical question, we offer a framework that makes genomic surveillance of carbapenem-resistant bacteria accessible and cost-efficient.

## Introduction

The rise of antimicrobial resistance—especially against last-line antibiotics such as carbapenems—has become an increasingly serious threat to global public health (1–3). Infections involving carbapenem-resistant *Enterobacterales* (CRE) leave physicians with few treatment alternatives and are commonly associated with delayed or inadequate care, higher morbidity, and increased mortality (3,4). Carbapenem resistance in *Enterobacterales* is predominantly disseminated via plasmid-borne carbapenemase genes, which can be transferred horizontally across bacterial species and lineages and thus limits the effectiveness of species-level surveillance for infection prevention and control (5–11). Consequently, surveillance approaches that go beyond the taxonomic identification of bacteria and instead incorporate plasmid-level resolution are required to understand and interrupt CRE persistence and transmission (11,12).

Currently established routine clinical diagnostics rely on species identification via matrix-assisted laser desorption ionisation time-of-flight mass spectrometry (MALDI-TOF MS), phenotypic antimicrobial susceptibility testing, and targeted carbapenemase detection assays. Although essential for guiding clinical decisions, these methods provide no information on the genomic context of resistance, such as on plasmid architecture—which is, however, required to confirm transmission events, cross-species dissemination, or atypical resistance mechanisms (5,13,14). Whole-genome sequencing (WGS), on the other hand, enables high-resolution characterisation of resistance determinants and their genomic context and has become an increasingly important tool in antimicrobial resistance surveillance. Nanopore sequencing technology generates highly accurate sequencing reads of any length, enabling complete *de novo* plasmid assemblies that traditional short-read sequencing approaches often cannot achieve (15–18). Nanopore sequencing can further be applied in real time at the point of care, with the potential to provide same-day whole-genome interpretation and to reduce costs by halting data generation as soon as the necessary sequencing depth is reached (13,19). Previous studies that benchmarked nanopore WGS against phenotypic carbapenem susceptibility tests relied on provisional nanopore sequencing chemistries with elevated raw-read error rates (R9.4.1 chemistry), limiting single-nucleotide and carbapenemase-subtype resolution (20). The R10.4.1 nanopore chemistry has been the first to achieve a highly accurate raw-read accuracy of roughly 99%, enabling highly accurate bacterial and plasmid genome assemblies from nanopore sequencing data alone (16,17). Using this chemistry, we showed that nanopore WGS can resolve plasmid-encoded carbapenem resistance reservoirs (11) as well as resistances to last-resort antibiotics (13)—which could not be resolved by established diagnostics.

Thus far, no systematic comparison of highly accurate nanopore WGS and established clinical diagnostics for CRE surveillance has been performed. Clear guidelines on necessary minimum sequencing depths are further required since insufficient data might compromise WGS data interpretation, potentially limiting the practical utility of nanopore WGS in clinical microbiology; this is especially relevant given the flexible data acquisition enabled by on-site nanopore sequencing. While previous research has suggested sequencing depths of 40× to obtain complete bacterial and plasmid genomes (16,17), robust threshold recommendations are needed for accurate CRE species identification, strain typing, carbapenemase detection, plasmid reconstruction, and plasmid-level epidemiology.

We here compared nanopore WGS and established diagnostics across 110 CRE isolates obtained from a tertiary-care hospital in Munich, Germany, to show the additional value of WGS for pathogen strain-level resolution, carbapenem resistance profiling, and plasmid epidemiology. In a subset of 100 carbapenemase-producing CRE isolates, we determined the sequencing depth required for accurate species identification, strain typing, carbapenemase detection, and plasmid-level epidemiology. Across all isolates, nanopore WGS correctly reproduced or superseded established diagnostics, and additionally resolved potential resistance mechanisms that established diagnostics had missed in a subset of ten CRE isolates.

## Materials and Methods

This study comprised 110 CRE isolates, namely 100 carbapenemase-producing *Enterobacterales* isolates (which tested positive for carbapenemase production by lateral flow tests) and 10 carbapenem-non-susceptible isolates (which showed phenotypic carbapenem non-susceptibility but tested negative for carbapenemase production by lateral flow tests), collected for routine diagnostics at a tertiary-care hospital in Munich, Germany, between 2022 and 2023. Most isolates (n=87) were obtained from routine carbapenem resistance surveillance via rectal or inguinal swabs; the remainder came from diagnostic samples including urine (n=8), soft tissue infection swabs (n=7), respiratory samples (n=6), catheter (n=1), and blood culture (n=1).

Ethical approval of this study was granted by the TUM ethics committee (2024-522-S-CB) in accordance with the principles of the Declaration of Helsinki. The requirement for informed patient consent was waived because all samples were collected as part of routine diagnostic and surveillance procedures; no patient-identifiable information is reported.

### Established diagnostics

Isolates were obtained from routine surveillance swabs or diagnostic samples (**Supplementary Table 1**). The samples were plated on BD® MacConkey (Becton Dickinson GmbH, Heidelberg, Germany) or Thermo Scientific™ Brilliance™ ESBL agar plates, then incubated at 37°C for 20-24 hours. Isolates were considered non-duplicates if they differed in species, carbapenemase type, or patient origin, or if sampling interval exceeded seven days. Species identification was performed from a single colony-forming unit (CFU) using MALDI-TOF MS (Bruker Daltonics GmbH, Bremen, Germany, MBT Compass 4.1, MBT Compass Reference Library 2023). Phenotypic antibiotic susceptibility testing was performed using VITEK 2 (BioMérieux, Marcy l’Etoile, France), with minimum inhibitory concentrations (MICs) being interpreted according to the European Committee on Antimicrobial Susceptibility Testing (EUCAST) guidelines (21). Carbapenemase production was assessed by a lateral flow immunochromatography assay consisting of a lateral flow assay (Resist-5 O.K.N.V.I panel, by CORIS BioConcept, Gembloux, Belgium) that detects KPC, OXA-48-like, NDM, VIM, IMP enzymes, according to the manufacturer’s instructions (11) (**Figure 1**).

**Figure 1.**
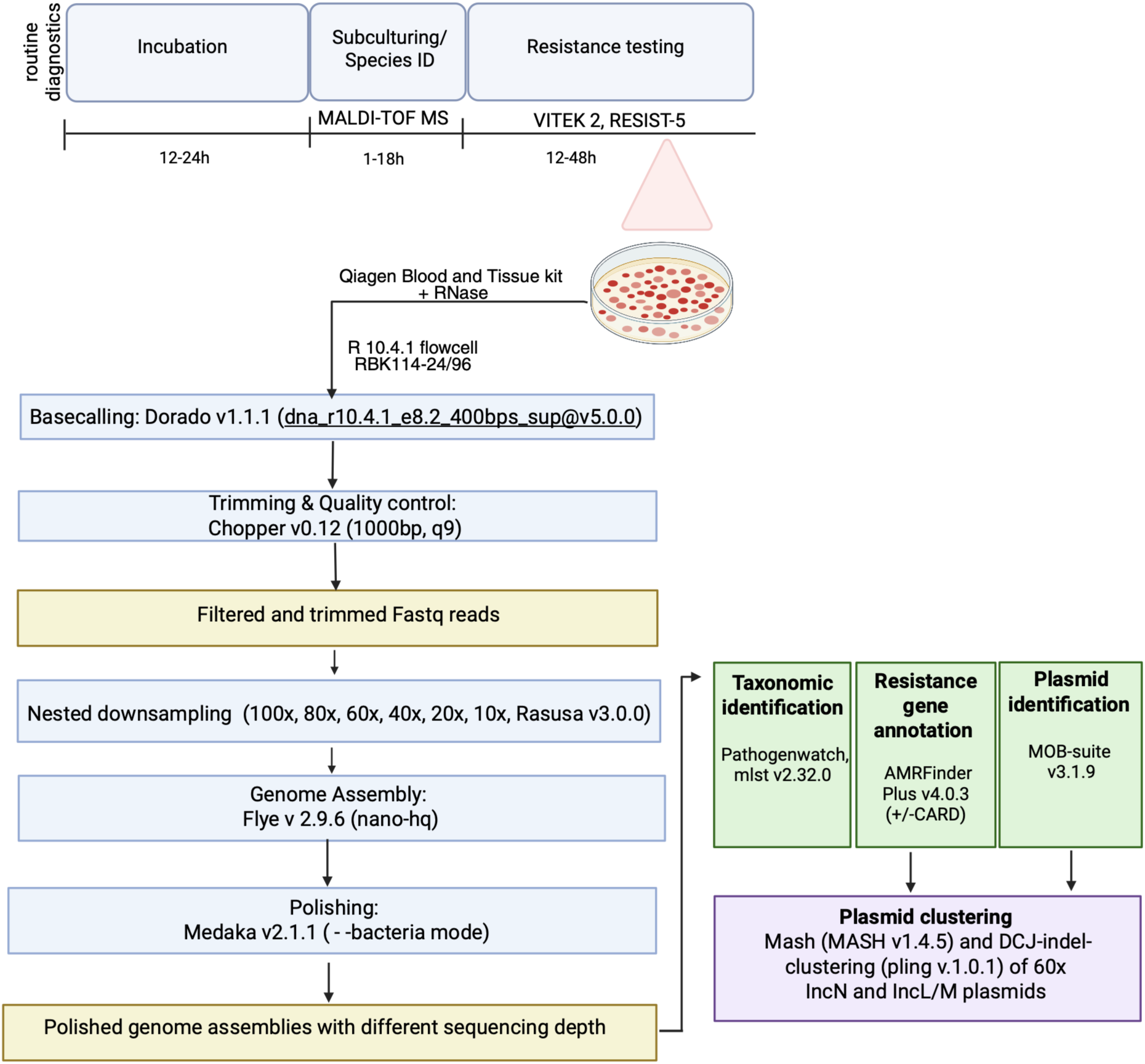
Overview of established diagnostics and nanopore sequencing-based WGS workflow. After incubation (12-24h) and subculturing, established diagnostics perform species identification by MALDI-TOF MS, phenotypic antimicrobial resistance testing (VITEK 2), and, for the isolates with elevated carbapenem minimum inhibitory concentrations (MICs), carbapenemase production is confirmed by lateral flow immunochromatography (RESIST-5) (Materials and Methods). The nanopore sequencing-based WGS workflow consists of DNA extraction including an RNase treatment step, library preparation, and sequencing. Raw nanopore sequencing reads are basecalled, trimmed, and filtered. Filtered reads are downsampled in a nested manner across a range of sequencing depths (10-100×), followed by *de novo* assembly and assembly polishing. Taxonomic annotation is conducted using Pathogenwatch for species-level resolution and the mlst tool for sequence typing. Assemblies are annotated for antimicrobial resistance genes, plasmid content, and replicon type. Carbapenemase-encoding plasmids are subjected to backbone-level and structural clustering (at 60× sequencing depth; Materials and Methods).

### DNA extraction and nanopore sequencing

DNA was extracted from overnight cultures using the Qiagen DNeasy Blood & Tissue kit (Gram-negative protocol) (22) with extended lysis (2h), 40 μL of proteinase K, and an RNase A treatment step to improve yield and purity, as described previously (11). Nanopore sequencing libraries were prepared using Oxford Nanopore Technologies’ SQK-RBK114-24/96 Rapid Barcoding Kit (two barcodes per sample) and sequenced on R10.4.1 flow cells for up to 72 hours on either a MinION Mk1d or PromethION 2 Solo sequencer, targeting a minimum mean sequencing depth of 40x (**Figure 1**).

### Nanopore sequencing data processing

All bioinformatic protocols are available at https://github.com/ElaSrbrn/AMR_clustering. Raw nanopore data were basecalled using Dorado v1.1.1 with the super-accuracy (SUP) model (dna_r10.4.1_e8.2_400bps_sup@v5.0.0). Adapter trimming and read filtering (Q<9; length <1 kbases (kb)) were performed with Chopper v0.12 (23). Sequencing summaries were generated using Seqkit v2.10.0 (24). Filtered reads were downsampled in a nested manner to defined target sequencing depths using Rasusa v3.0.0 (seed = 42) (25), assuming a conservative mean *Enterobacterales* genome size of 5 Mb; total sequencing depth was defined as the total number of sequenced bases divided by this assumed genome size. For samples with initial sequencing depth ≥ 100×, reads were first downsampled to 100× and then in a nested manner to 80×, 60×, 40×, 20×, and 10×; for samples with lower initial sequencing depth, reads were downsampled only to the highest achievable depth and its nested subsets (**Figure 1**).

### *De novo* assembly and annotation

*De novo* assemblies were generated using Flye v2.9.6 (26,27) for high-quality long reads (nano-hq mode) and polished with Medaka v2.1.1 (--bacteria mode, https://github.com/nanoporetech/medaka, accessed on 22 July 2025). Sequencing reads were aligned to their respective polished assemblies using minimap2 (28). Assembly completeness and contamination were evaluated on polished assemblies using CheckM2 v1.1.0 (29) (**Figure 1**). For a single isolate, a consensus assembly was generated with Autocycler v0.5.2 at a sequencing depth of 40x, based on outputs from the assembly tools Raven, myloasm, miniasm, Flye, metaMDBG, NECAT, NextDenovo and Plassembler.

Species identification was performed using Pathogenwatch v2.3.1 (30). Multi-locus sequence typing (MLST) was performed to obtain sequence types (STs) using the mlst tool v2.32.0 (31). Antimicrobial resistance genes were identified using AMRFinderPlus v4.0.3 (32–34). Carbapenemase-encoding contigs were functionally annotated with MOB-suite v3.1.8 (35) to assign replicon types, relaxase families, mating-pair formation systems, and predicted mobility status. In some carbapenem-non-susceptible isolates in which no carbapenemase gene was detected, additional screening was conducted (*i*) using the Comprehensive Antibiotic Resistance Database (CARD) via ProkSee.com (DB v4.0.1, accessed 5 May 2026) (**Figure 1**) (36,37), and (*ii*) using Bakta v1.11.4 for gene annotation followed by point mutation calling using AMRFinderPlus v4.0.3 while specifying each isolate’s species (--organism) to identify potential regulatory mutations associated with reduced porin expression.

At a sequencing depth of 20x, assembly-based carbapenemase gene detection was compared with sequencing read-level detection using AMRFinderPlus as previously described. Single-read carbapenemase detections were discarded if coverage or identity were below 95%.

### Plasmid reconstruction analyses

To evaluate the effect of sequencing depth on plasmid reconstruction, the contig size of each carbapenemase-encoding plasmid at 10×, 20×, and 40× was compared with the size of the same carbapenemase-encoding plasmid in the respective 60× assembly. The 60× assembly served as the reference since this sequencing depth was attainable for most isolates (98%) and gave the broadest coverage of isolates and plasmid types. A plasmid was classified as stably reconstructed at a sequencing depth when its contig size deviated by no more than 5% in either direction from its 60× size. It was classified as unstable when the size deviation exceeded 5%, or the plasmid replicon was not typable. Sensitivity analyses were repeated at deviation thresholds of 3% and 10%.

### Plasmid similarity analyses

Plasmid clustering was performed at 60× sequencing depth for the two most abundant carbapenemase-encoding replicon types (IncN, IncL/M). Pairwise genomic distances between plasmids of the same replicon types were first estimated using MinHash-based approximation of Jaccard distances with MASH v2.3 (38) with default parameters (sketch size 1,000 and k-mer size 21). Plasmids were retained for downstream analysis if they had a lenient Mash distance of ≤0.005 with at least one other plasmid of the same replicon type, indicating a conserved backbone (39) rather than near-identity, which would be indicative of recent transmission and require a stricter threshold of typically 0.001 (11). Retained plasmids were then analysed with pling v1.0.1 (40) and assigned to communities at a containment threshold of <0.3 and to subcommunities at a double-cut-and-join (DCJ)-Indel distance of ≤4, clustering plasmids that are structurally consistent with recent common ancestry or transmission (40).

### Statistical analyses

All statistical analyses were conducted in R v4.3.1, using tidyverse suite (v2.0.0; dplyr v1.1.4, ggplot2 v3.5.2) and pheatmap v1.0.13 for visualisations. For assessing carbapenemase-encoding plasmid reconstruction across sequencing depths, the association between sequencing depth and plasmid reconstruction was modelled by generalised estimation equations (GEE) to account for repeated measurements per plasmid (across sequencing depths) (geepack v1.3.12) (41) and confirmed with Cochran’s Q test.

## Results

### Established diagnostics

Across the 100 carbapenemase-producing *Enterobacterales* isolates, the three most frequent species were *Klebsiella pneumoniae* complex (n=32), *Escherichia coli* (n=22) and *Citrobacter freundii* complex (n=19) according to MALDI-TOF MS (Materials and Methods; **Supplementary Table 1**; **Figure 2**). Carbapenem resistance was confirmed across all isolates using VITEK 2 (Materials and Methods). In total, 105 carbapenemases were detected across the 100 isolates using the lateral-flow assay RESIST-5 (Materials and Methods), with five isolates carrying two carbapenemase genes, respectively: Four isolates combined an OXA-48-like enzyme with a metallo-β-lactamases (VIM or NDM), and one isolate co-produced OXA-48-like and KPC (**Supplementary Table 1**; **Figure 2**). Carbapenemase distribution varied by species: KPC and OXA-48-like enzymes predominated in *K. pneumoniae* complex isolates; *E. coli* showed a broader distribution, including OXA-48-like, NDM, and KPC; and *C. freundii* complex isolates were primarily associated with OXA-48-like enzymes (**Supplementary Table 1**; **Figure 2**).

**Figure 2.**
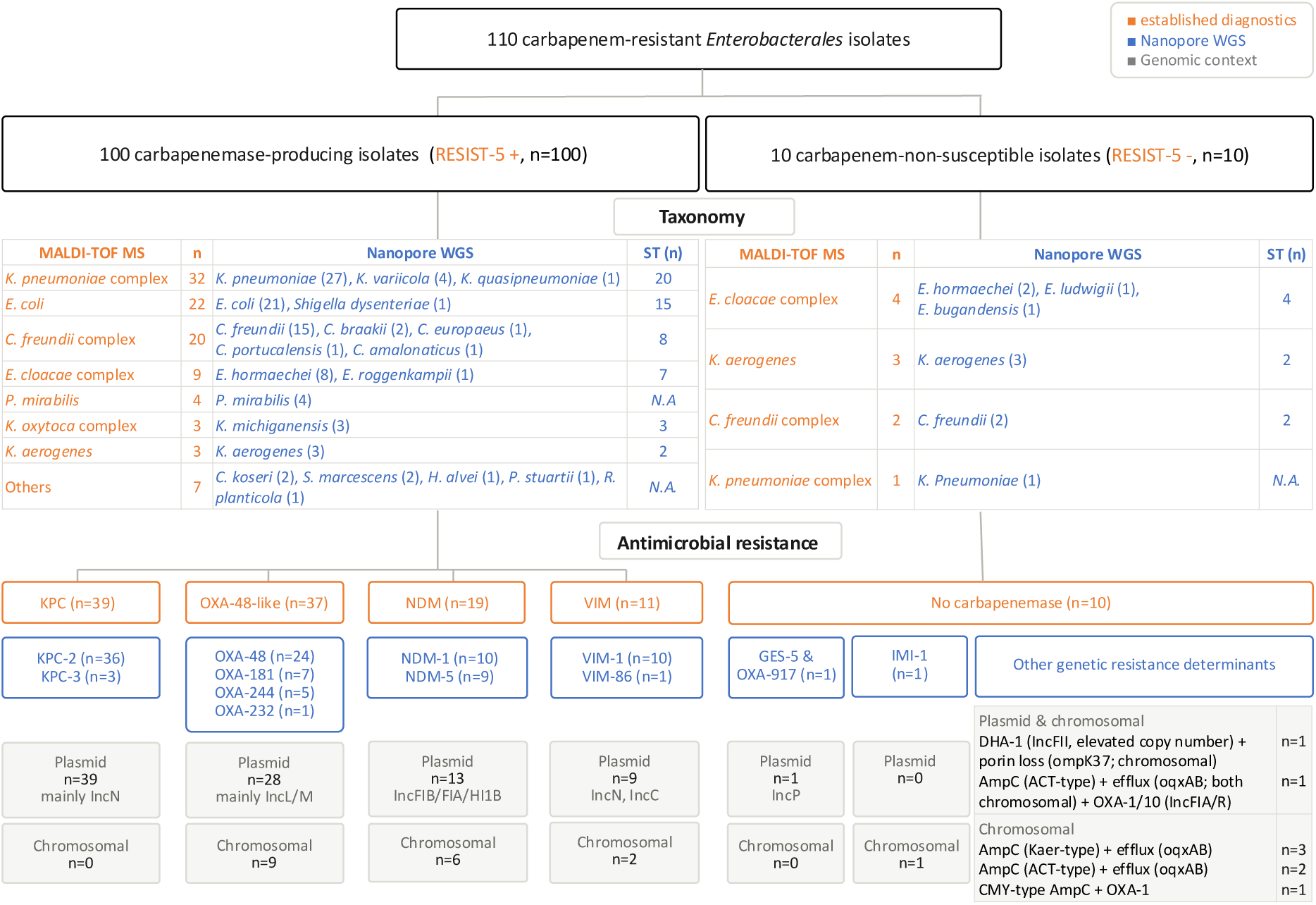
Comparison of nanopore whole-genome sequencing (WGS) and established diagnostics across 110 carbapenem-resistant (100 carbapenemase-producing and 10 carbapenem-non-susceptible) *Enterobacterales* isolates. Established diagnostics consist of MALDI-TOF MS for species (or species complex) identification, VITEK 2 for phenotypic carbapenem resistance prediction, and RESIST-5 lateral flow assays to confirm the presence of carbapenemase gene families; isolates with phenotypic carbapenem resistance but negative carbapenemase gene family detections are classified as carbapenem-non-susceptible. Nanopore WGS assemblies are annotated for bacterial species and, where applicable, sequence type (ST) identification, carbapenemase subtype detection, genomic location of the carbapenemase on a plasmid or the bacterial chromosome, and, where applicable, replicon type of the plasmid (**Supplementary Table 1**; Materials and Methods).

The additional 10 *Enterobacterales* isolates that showed phenotypic carbapenem non-susceptibility using VITEK 2 but tested negative for carbapenemase production by the RESIST-5 assay, consisted of 4 *E. cloacae* complex, 2 *C. freundii* complex, 3 *K. aerogenes* and 1 *K. pneumoniae* complex isolate according to MALDI-TOF MS. Here, established diagnostics could not account for the observed resistance phenotype (**Supplementary Table 1**; **Figure 2**).

### Genomic data

Across all 110 isolates, nanopore sequencing generated 123.9 Gb and 18.6M sequencing reads. After read filtering, the isolate median values were 708 Mb (interquartile range (IQR): 445–1,149 Mb), 109,400 reads (IQR: 66,100–185,000), N50 of 10.4 kb (IQR: 9.0–13.8 kb), 54.2% for GC content (IQR: 50.7–56.2%), and 19.6 for mean quality (IQR: 18.3–21.9), with a median 91% of bases at ≥Q20 (IQR: 87–94%) and 84% at ≥Q30 (IQR: 80–89%).

Depending on the sequencing throughput, a different subset of the 100 carbapenemase-producing *Enterobacterales* isolates could be analysed across sequencing depths: All 100 carbapenemase-producing isolates could be analysed at 40×, 98 at 60×, 76 at 80×, and 62 at 100x. The summary of the WGS nanopore results in **Supplementary Table 1** is based on a sequencing depth of 40x, which could be achieved for all isolates. The sequencing throughput was independent of the isolate’s bacterial species or carbapenemase family; downsampling to different sequencing depths did not affect read length distribution and quality metrics (Materials and Methods)

### Carbapenemase-producing *Enterohbacterales* isolates

#### Taxonomic identification

At a low sequencing depth of 10x, established diagnostics and nanopore WGS agreed on the species identification for all 100 carbapenemase-producing isolates, considering that *E. coli* and *Shigella* species cannot be reliably differentiated by MALDI-TOF MS (**Figure 2**; **Supplementary Table 1**; Materials and Methods). For 64 isolates, MALDI-TOF MS only reported a species complex while nanopore WGS was invariably able to resolve the taxonomic classification to the species level (**Figure 2**; **Supplementary Table 1**).

MLST assignment was stable across sequencing depths for the 81 typable carbapenemase-producing isolates (Materials and Methods). Only at a depth of 10x, two *E. coli* isolates resulted in no ST since the wrong *Salmonella enterica* scheme was selected; the ST of these two isolates could be recovered at a depth of 20x (**Figure 2**; **Supplementary Table 1**). Of the 19 isolates without a ST, eight isolates belonged to species with no established MLST scheme (*Proteus mirabilis*, n=4; *Serratia marcescens*, n=2; P*ovidencia stuartii*, n=1; and *Hafnia alvei*, n=1), six isolates returned inexact or partial allele calls *Citrobacter* spp., n=1; *E. coli*, n=1; and *Raoultella planticola*, n=1), four isolates returned a complete profile of exact allele matches for which the MLST reference database contained no corresponding sequence type (*K. pneumoniae*, n=1; *K. variicola*, n=2; and *K. aerogenes*, n=1), and one isolate matched the *C. freundii* ST22 profile except for a duplicated *lysP* locus.

#### Carbapenemase gene detection and genomic context

In total, nanopore WGS detected 106 carbapenemase genes across the 100 isolates (Materials and Methods). Carbapenemase detection was robust across sequencing depths, with most genes identified at a depth of 10×; missed genes (n=2) and missed subtypes (n=5) could be reliably recovered at a depth of 20x—except for a single *bla*_KPC-3_ gene, which was located on a fragmented IncFIB/IncFII plasmid and required a sequencing depth of 60x (**Supplementary Table 2**). The *bla*_KPC-3_ gene could, however, be recovered at a sequencing depth of 20x when antimicrobial resistance gene annotation was applied to individual sequencing reads; in general, read-based analysis accurately recovered all carbapenemase genes, but only do the gene family instead of subtype level in the case of seven carbapenemase detections (Materials and Methods).

Carbapenemase genes were identified in chromosomal and plasmid contexts. *bla*_KPC-2_ (n=36), and *bla*_OXA-48_ (n=24) were exclusively plasmid-encoded, whereas *bla*_OXA-244_ (n=5) was exclusively chromosomally encoded. *bla*_NDM-5_, *bla*_NDM-1_, *bla*_OXA-181_, and *bla*_VIM-1_, were identified in both contexts (**Supplementary Table 1**). Chromosomal copies of *bla*_OXA-181_ were annotated by MOB-suite as harbouring IncX3-associated replicon sequences, consistent with prior integration from an IncX3 plasmid (42). Distinct associations were observed between carbapenemase subtypes and plasmid replicon subtypes (**Figure 3**). *bla*_KPC-2_ was predominantly carried on IncN plasmids, and *bla*_OXA-48_ on IncL/M plasmids. *bla*_NDM-5_ and *bla*_KPC-3_ were exclusively associated with IncF plasmids, while *bla*_VIM-1_ showed a broader distribution across multiple plasmid types (**Supplementary Table 2**; **Figure 3**).

**Figure 3.**
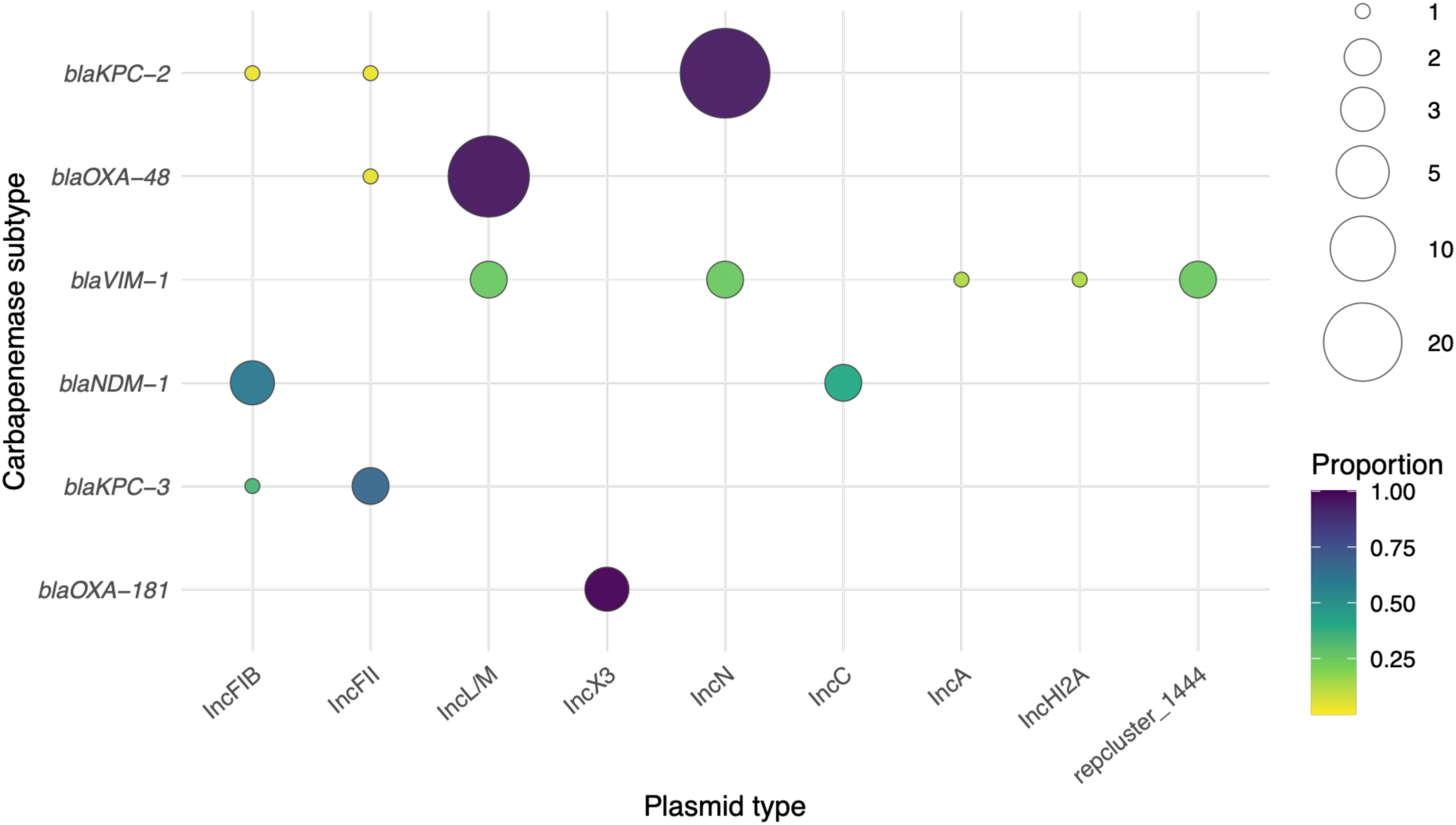
Plasmid-borne carbapenemase subtypes (y-axis) across main plasmid replicon types (x-axis) detected in 100 carbapenemase-producing *Enterobacterales* isolates. The bubble size represents the number of detections, and the colour scheme represents the proportion of each plasmid type within a given carbapenemase subtype. Plasmids were assigned a single replicon class per isolate to avoid overcounting composite plasmids. Specific replicon types were assigned only when no Inc-type replicon was detected. Carbapenemase subtypes *bla*VIM-86, *bla*NDM-5 and *bla*OXA-232 were excluded as they were only detected on one plasmid type, respectively (Materials and Methods).

#### Plasmid reconstruction and epidemiology

Plasmid reconstruction stability was assessed by comparing the contig size of each of altogether 86 carbapenemase-encoding plasmids at 10×, 20×, and 40× with its 60× reference assembly (deviation threshold of 5%; Materials and Methods; **Supplementary Table 3**). For 9 of the 86 plasmids (10.5%), the 60× reference contig was not circularised. Across sequencing depths, 78% of the plasmids were stably reconstructed at 10× (67/86), increasing to 90% at 20× (77/86), and plateauing at 92% at 40× (79/86). The increase in the number of stably reconstructed plasmids with sequencing depth was significant (GEE p = 0.008; Cochran’s Q p = 0.002) and driven primarily by an increase in sequencing depth to 20x. Stability classifications were robust across alternative deviation thresholds of 3% and 10% (GEE trend p = 0.006 and 0.007, respectively; **Supplementary Table 3**). At 80×, 92% of assessable plasmids were stably reconstructed (57/62), matching the plateau reached at 40x. A single small carbapenemase-encoding plasmid (isolate 6737, *bla*_OXA-232_, <10 kb) was reconstructed inconsistently across all depths using our computational assembly pipeline; however, the implementation of the consensus assembly tool Autocycler generated a stable circular plasmid already at a sequencing depth of 40x (Materials and Methods).

The most common carbapenemase-encoding plasmids across our 100 carbapenemase-encoding Enterobacterales isolates were IncN (n=35) and IncL/M (n=23). Of 35 carbapenemase-encoding IncN plasmids, 33 plasmids (94%) shared a pairwise Mash distance of ≤0.005 with at least one other IncN plasmid and were retained for downstream clustering. For the 23 IncL/M plasmids, this applied to 22 plasmids (96%) (**Supplementary Table 4**; Materials and Methods). All retained IncN plasmids grouped into a single subcommunity based on DCJ-indel distance metrics, indicating a broadly conserved backbone structure (**Figure 4A**; **Supplementary Table 5**). The retained IncL/M plasmids showed considerably greater structural diversity, with pairwise DCJ-indel distances ranging from 0 to 10 (**Figure 4B**; **Supplementary Table 5**).

**Figure 4.**
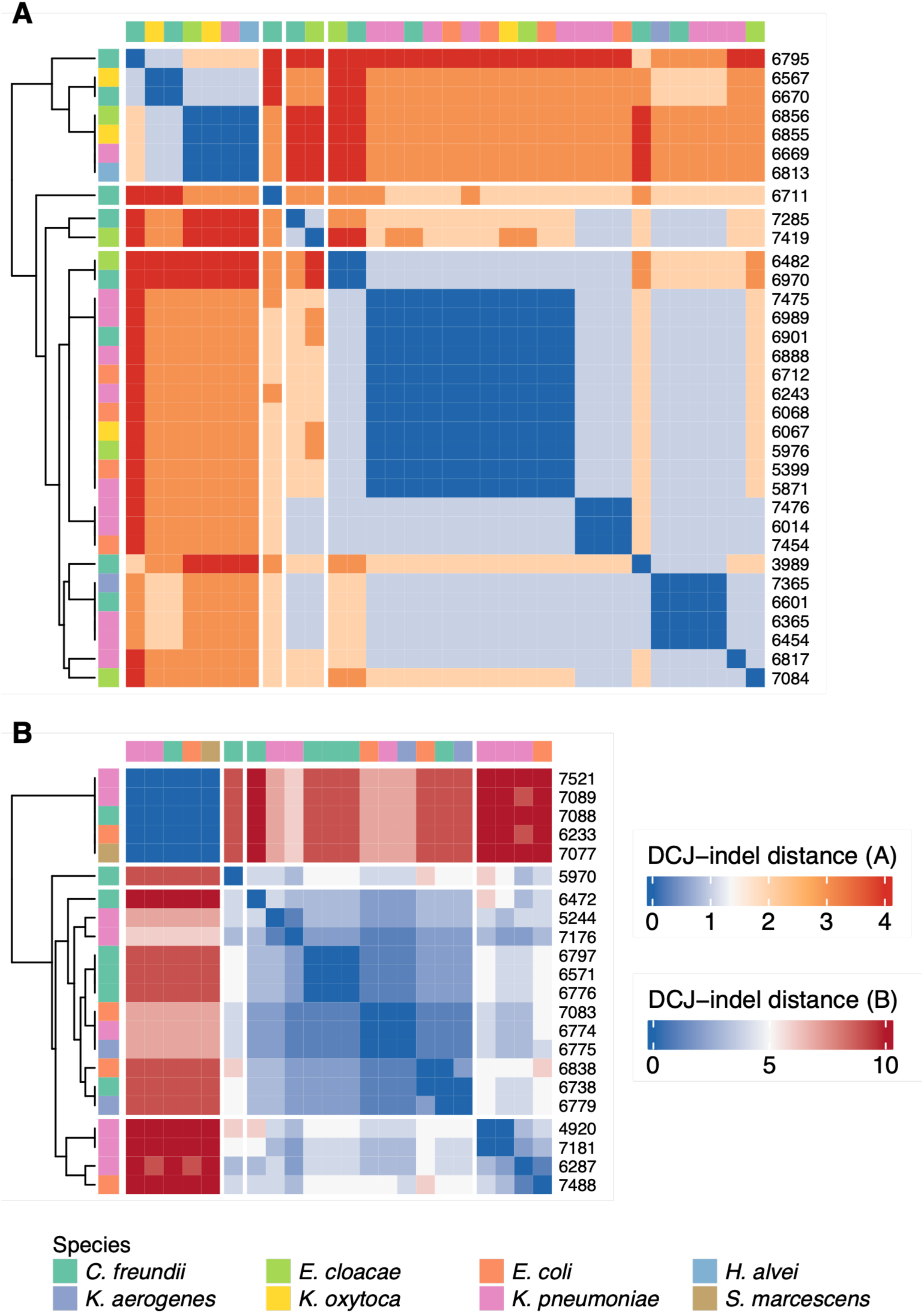
Pairwise clustering of carbapenemase-encoding plasmids. Hierarchical clustering heatmap of pairwise DCJ–indel distances of IncN or IncL/M plasmids after Mash distance filtering (≤0.005). The colour scheme represents DCJ-indel distances from 0 (blue; structurally identical) to the respective largest distance (red; most divergent). Row and column colour bars indicate bacterial species (see **Supplementary Table 1** for isolate details). **A.** Hierarchical clustering of 33 IncN plasmids. **B.** Hierarchical clustering of 22 IncL/M plasmids.

### Carbapenem-non-susceptible *Enterobacterales*

As in the case of the carbapenemase-producing *Enterobacterales* isolates, nanopore WGS at least matched the taxonomic resolution of established diagnostics across all ten carbapenem-non-susceptible *Enterobacterales* isolates. The four *E. cloacae* complex isolates were resolved to three species, *E. hormaechei* (n=2), *E. ludwigii* (n=1) and *E. bugandensis* (n=1), the two *C. freundii* complex isolates were both resolved to *C. freundii*, and the one *K. pneumoniae* complex isolate could be resolved to *K. pneumoniae*. Nine of the ten isolates could additionally be assigned to a ST; the *K. pneumoniae* isolate did not match any reference ST while returning a complete profile of exact allele matches (**Figure 2**; **Supplementary Table 1**).

In two of the ten carbapenem-non-susceptible isolates, WGS assemblies detected the relevant carbapenemase genes (Materials and Methods). One *C. freundii* isolate (ST19) carried two plasmid-associated carbapenemase genes, *bla*_GES-5_ and *bla*_OXA-917_ (an OXA-427-like enzyme (43)), on an IncP plasmid, and one *E. bugandensis* isolate (ST901) carried the chromosomally encoded carbapenemase *bla*_IMI-1_ (**Supplementary Table 1**; **Figure 2**).

In the remaining eight isolates, no carbapenemase was detected by WGS. Six isolates carried intrinsic chromosomally encoded AmpC β-lactamases and oqxAB efflux pumps: three *K. aerogenes* (AmpC-Kaer), two *E. hormaechei* and one *E. ludwigii* (ACT-type) isolates, with one of them additionally carrying *bla*_OXA-1_ and *bla*_OXA-10_ on IncFIA and IncR plasmids. One *C. freundii* isolate carried an intrinsic CMY-type AmpC β-lactamase as well as *bla*_OXA-1_, and one *K. pneumoniae* isolate carried the AmpC β-lactamase *bla*_DHA-1_ (44) on a plasmid contig with approximately sevenfold higher sequencing coverage than the bacterial chromosome (45,46) (**Supplementary Table 1**; **Figure 2**). Additional analyses identified genetic signatures of porin loss or downregulation, with *OmpA* and *OmpK37* variants (47,48) in the *K. pneumoniae* isolate (isolate 7320) and *ramR* loss-of-function mutations in the same *K. pneumoniae* isolate and in one of the *K. aerogenes* (5861) isolates (**Supplementary Table 1**; Materials and Methods).

## Discussion

Bacterial whole-genome reconstruction by nanopore sequencing has been proposed as a rapid high-resolution complement to established clinical diagnostics when it comes to pathogen and antimicrobial resistance profiling (11–13,15–17,19,49). While on-site nanopore whole-genome sequencing (WGS) thus promises to be especially relevant for monitoring carbapenem-resistant *Enterobacterales* (CRE), whose rapid plasmid-mediated dissemination of carbapenemase genes pose a critical public health threat (2,6,10,11,50), no systematic comparison between highly accurate (R10.4.1 chemistry) nanopore WGS and established diagnostics has been conducted to date.

We used 110 carbapenemase-producing and carbapenem-non-susceptible *Enterobacterales* isolates from routine diagnostics to directly compare established diagnostics and nanopore WGS in terms of species identification, strain typing, carbapenemase detection, and plasmid-level epidemiology, and determined minimum sequencing depth requirements for each of these clinically relevant tasks. Across the carbapenemase-producing isolates, nanopore WGS reproduced routine carbapenemase family and pathogen detections, and additionally resolved carbapenemase subtypes and genomic contexts as well as bacterial species and sequence type (ST) beyond what established diagnostics could achieve. For CRE surveillance and infection prevention and control, this information on the bacterial ST, the carbapenemase subtype, and the carbapenemase-carrying plasmid can help distinguish clonal spread from plasmid dissemination—a distinction that established diagnostics can usually not make (11,12). Across the carbapenem-non-susceptible isolates, for which established diagnostics confirmed phenotypic carbapenem non-susceptibility but could not detect the relevant carbapenemase gene, nanopore WGS identified potential resistance mechanisms that established diagnostics had missed. Our results thus strongly support the integration of nanopore WGS into CRE diagnostics, surveillance, and outbreak investigation.

From a clinical microbiology perspective, the implementation of nanopore WGS into routine diagnostics and surveillance requires clear guidelines on required sequencing throughput—to minimise costs, speed up *de novo* genome generation, and avoid suboptimal interpretation due to insufficient data (51–55). We here determined the minimum sequencing depths required for accurate CRE species identification, strain typing, carbapenemase detection, and plasmid-level epidemiology. A low sequencing depth of 10× was sufficient for reliable species identification: Considering that *E. coli* and *Shigella* species cannot be reliably differentiated by MALDI-TOF MS (56), taxonomic identification was concordant with established diagnostics across all 110 isolates while resolving the taxonomic identification of 71 of these isolates, for which MALDI-TOF MS only reported a species complex, to the species level. Subsequent MLST assignment was feasible for 90 of the 110 CRE isolates, with robust annotations across all sequencing depths. Only at a sequencing depth of 10x, two *E. coli* isolates were not assigned a correct ST; we thus recommend a minimum sequencing depth of 20x for reliable MLST assignments of CRE isolates. The remaining 20 isolates either belonged to bacterial species without MLST schemes or only returned partial reference matches of complete allelic profiles; these dropouts therefore reflect database coverage and locus-level ambiguity rather than insufficient sequencing depth.

Across the 100 carbapenemase-encoding *Enterobacterales* isolates, nanopore WGS detected 106 carbapenemase gene subtypes, thus recovering all but one carbapenemase predicted by the established RESIST-5 assay at a sequencing depth of 20x. The one missed carbapenemase, a single *bla*_KPC-3_ gene, could only be recovered at 60x. While the RESIST-5 assay only reports the carbapenemase family (i.e., KPC, OXA-48-like, NDM, VIM, or IMP), nanopore WGS was able to identify all carbapenemase subtypes, separating epidemiologically and functionally distinct carbapenemases, such as *bla*_KPC-2_ from *bla*_KPC-3_, or *bla*_OXA-48_ from *bla*_OXA-181_, *bla*_OXA-232_, and *bla*_OXA-244_, and determined whether the carbapenemase was plasmid-borne or chromosomally encoded. As we hypothesised that the *bla*_KPC-3_ gene was missed due to the fragmented nature of the respective IncFIB/IncFII plasmid assembly, we followed up with carbapenemase detection analyses on the sequencing read level, where the *bla*_KPC-3_ gene could indeed be recovered—even at a relatively low sequencing depth of 20x. In general, all carbapenemases could be detected without assembly at this sequencing depth, albeit seven of them only on the gene family level. The comparison of assembly- and read-level carbapenemase detections suggests that a combination of both approaches might enable the precise identification of carbapenemases and their genomic context (assembly-level) as well as the rapid detection of carbapenemases or carbapenemase families (read-level), especially in the case of problematic and fragmented *de novo* assemblies.

A higher sequencing depth of roughly 40x was required for plasmid-level epidemiological inferences, which allowed for stable reconstruction of plasmids in 92% of the isolates. A single small (<10 kb) *bla*_OXA-232_-encoding plasmid was inconsistently reconstructed across all sequencing depths. The unstable reconstruction of this small plasmid was due to the implementation of the assembly tool Flye in our computational pipeline, which is known to be limited for the reconstruction of small assemblies (15,57). While the highly optimised consensus-based assembly tool Autocycler (58) successfully reconstructed this small plasmid at a sequencing depth of 40x, it is computationally expensive and its application might not be feasible at the point of care and without the usage of high-performance compute. To the contrary, our entire workflow can be performed on a mobile platform, with data analysis being feasible on a high-performance laptop, supporting implementation across a range of clinical settings including resource-limited environments (13). We hope that our recommendations on sequencing throughput and computational analysis thus enable the cost-efficient and possibly on-site integration of nanopore WGS into CRE diagnostics, surveillance, and outbreak investigation.

In ten carbapenem-non-susceptible CRE isolates, established diagnostics did not find the relevant carbapenemase gene despite phenotypic evidence of resistance. In two of these isolates, nanopore WGS could recover the relevant carbapenemase, a plasmid-borne *bla*_GES-5_ together with an OXA-427-like enzyme in a *C. freundii* isolate and a chromosomally encoded IMI-type enzyme in one *E. bugandensis* isolate. As the RESIST-5 assay is restricted to other carbapenemase families, the negative RESIST-5 results in these two isolates make sense and show that negative RESIST-5 results cannot reliably exclude carbapenemase production (59). Together with the limitation of the resolution of RESIST-5 to the gene family level, our findings illustrate the added value of whole-genome data in comparison to targeted molecular panels (13,59,60).

In the remaining eight carbapenem-non-susceptible isolates, nanopore WGS detected AmpC β-lactamases, which are known to raise carbapenem minimum inhibitory concentrations (MICs) when hyperproduced in combination with reduced outer membrane permeability. In seven isolates, AmpC genes were intrinsic and chromosomally encoded, and accompanied by the presence of potentially relevant efflux pumps and/or OXA genes. One *K. pneumoniae* isolate carried the plasmid-borne AmpC *bla*_DHA-1_ with an elevated plasmid copy number of roughly sevenfold, potentially consistent with elevated plasmid copy number as a non-carbapenemase-mediated contributor to carbapenem resistance (45). The same isolate also carried *OmpA* and *OmpK37* variants, and, together with one other *K. aerogenes* with an intrinsic AmpC gene, also *ramR* loss-of-function mutations. The loss of *ramR* function might lead to *ramA* overexpression and thus repress major porin transcription. These mutations might point towards the role of reduced outer membrane permeability through porin disruption or similar mechanisms (14,61). To, however, quantify individual contributions and interactive effects of more complex genetic mechanisms of phenotypic carbapenem susceptibility, larger-scale systematic assessments of different genetically encoded combinations of enzymatic and permeability-based mechanisms in combination with phenotypic and functional validation will be required. Our study is limited due to the very small sample size of carbapenem-non-susceptible isolates (n=10), and due to retrospective instead of prospective inferences, which allowed us to assume carbapenem resistance across all isolates.

In our specific dataset, the genetic mechanisms of carbapenem resistance were predominantly plasmid-borne, which is consistent with the established role of horizontal gene transfer in the dissemination of carbapenem resistance and highlights the importance of plasmid-level instead of species-level surveillance of carbapenem resistances (5,8,9,62). Bacterial isolates of the same species and ST frequently carried different carbapenemases on different plasmid replicon types, ruling out clonal spread as the main driver of carbapenem resistance. Epidemiological analyses of the two main plasmid replicon types revealed a highly conserved IncN plasmid backbone, with clusters spanning several bacterial species and months of sampling in line with known interspecies persistence of a shared plasmid backbone within the hospital setting (11,63), while the IncL/M plasmids exhibited greater structural heterogeneity, resulting in multiple distinct clusters (64,65).

In summary, we show across 110 clinical isolates with phenotypic carbapenem resistance that nanopore WGS can at least reproduce and often supersede established diagnostics in terms of pathogen strain-level resolution, carbapenem resistance profiling, and plasmid epidemiology. Our findings thus support the integration of nanopore sequencing into routine clinical diagnostics, surveillance, and outbreak investigation. By providing laboratory and computational workflows as well as guidance on minimum sequencing throughput, we hope to enable the rapid and cost-efficient implementation of on-site genomics at the point of care.

## Data availability

The genomic dataset is available at ENA at project number PRJEB123044. All other supporting data are provided in the article and supplementary data files. All bioinformatic protocols are available at https://github.com/ElaSrbrn/AMR_clustering.

## Author contributions

ES and LU designed the DNA extraction and sequencing protocols. ES extracted and sequenced the DNA. KS conducted susceptibility testing, interpretation and species identification of query isolates. ES, SV and AS conducted the bioinformatic analysis under the supervision of LU and EFN. FG and NW provided samples, laboratory equipment and clinical expertise. LU, ES and EFN wrote the manuscript with input from all co-authors. LU supervised and financed the study.

## Conflicts of interest

LU has received travel expenses from Oxford Nanopore Technologies to present preliminary results of this study.

## Funding information

This project was funded by a Helmholtz Principal Investigator Grant and the Assistant Professorship “One Health with focus on microbial genomics and AI” (Institute for Food Safety and Hygiene, University of Zurich, Switzerland) awarded to LU. Laboratory and staff costs for routine diagnostics were covered by the Technical University of Munich Hospital and the Institute of Medical Microbiology, Immunology and Hygiene.

## Acknowledgements

We thank all laboratory technicians and physicians, especially Hannah Zierer and Madlen Peisker, at the Technical University of Munich Institute of Medical Microbiology, Immunology and Hygiene for their support. Figure 1 was generated using Biorender.

## Notes

### Summary of Updates

Typos and unclear statements corrected. A previously suggested mismatch between sequencing and established diagnostics was resolved due to a sample mix-up in established diagnostics. Read-level anaylses were added in addition to assembly-based analyses.

